# Seizures drive tau propagation in a tauopathy mouse model

**DOI:** 10.64898/2026.03.14.711088

**Authors:** Aaron J. Barbour, Keegan Hoag, Virginia M.Y. Lee, Delia M. Talos, Frances E. Jensen

**Author notes:** Corresponding authors: Frances E. Jensen, MD, FACP Arthur Knight Asbury MD Professor, Chair, Department of Neurology, Hospital of the University of Pennsylvania, 3400 Spruce Street, Dulles 3, Philadelphia, PA 19104, PA USA, Delia Talos, MD, 415 Curie Boulevard, 263 Clinical Research Building, Philadelphia, PA, 19104.

## Abstract

A bidirectional relationship between seizures and neurodegenerative disease has been established with neurodegenerative pathology found in late-onset epilepsy patients, increased risk of seizures in tauopathies, and accelerated Alzheimer’s disease progression in patients with epileptiform activity. Tau pathology spreads between interconnected neuronal networks, driving disease progression. We hypothesized that seizures would promote tau propagation throughout the brain in a tauopathy mouse model. To explore the brain-wide relationship between tau pathology and seizure activity, we crossed the T40PL-GFP mouse, which contains a pathogenic *MAPT* mutation tagged with GFP, with targeted recombination in active population (TRAP; T40PL-TRAP) mice to label all seizure activated neurons with tdTomato. We triggered tau propagation in these mice with intracerebral seeding of human AD brain-derived tau lysate and induced seizures with pentylenetetrazol (PTZ) kindling. With light sheet microscopy, we imaged and mapped tau-GFP and tdT levels throughout whole brain. We found that PTZ induced seizures worsened tau pathology in brain regions with increased tdT levels, including the hippocampus and cortex, and in the fiber tracts in T40PL-TRAP mice. We also found that seizure-activated (tdT+) neurons were more likely to develop somatic tau pathology compared to the surrounding (tdT-) populations. Overall, these data demonstrate that seizures can enhance tau pathology propagation.

## Introduction

The tau protein is expressed ubiquitously in neurons and normally functions to stabilize axonal microtubules (1). However, in pathological conditions tau can become hyperphosphorylated, leading to the mislocalization and aggregation of tau protein, leading to the formation of insoluble aggregates and neurofibrillary tangles (2). Mutations to the tau encoding *MAPT* gene, including proline 301 to leucine or serine (P301L, P301S), destabilizes the tau protein and is associated with familial tauopathies (3-6). The formation and spread of tau pathology occurs through interconnected neuronal networks, and is central to the progression of several neurodegenerative disorders, including Alzheimer’s disease (AD), frontotemporal lobar dementia with tau pathology (FTLD-tau), and progressive nuclear palsy (PSP) (7-11), but has also been seen to accumulate in other conditions such as epilepsy and TBI (12-14). With respect to AD, tau propagation throughout the brain can be modelled via intracerebral injection of pathological tau, including tau protein isolated from AD brain tissue (AD-tau) (15-18). Several mechanisms likely underlie the spread of tau pathology along neuroanatomical connections. We recently demonstrated a vulnerability of neuronal networks that displayed early neuronal hyperactivation to tau spread in an AD mouse model (18). Indeed, mounting evidence suggests that a primary mechanism underlying neuron-to-neuron tau spread is driven by synaptic transmission (7, 19-21). Given that tau pathology is central to multiple neurodegenerative disorders, understanding the networks impacted and the factors that drive tau propagation is critical to identify and develop therapies to slow disease progression across a range of disorders.

Seizures are a common comorbidity in tauopathies such as AD and FTLD (22-24). Subclinical epileptiform activity is found in >50% of AD patients (25, 26) and P301S *MAPT* mutation in humans causes a hereditary early onset version of FTLD with seizures (6). Furthermore, a bidirectional relationship between tau pathology and seizures has been established, as temporal lobe epilepsy patients display tau pathology in the hippocampus and temporal cortex (12, 13). In addition, prior studies have demonstrated increased seizure susceptibility in response to chemoconvulsants in tauopathy rodent models (27, 28). Notably, tau knockdown reduced seizure severity in genetic AD and epilepsy models and WT mice (29-33), demonstrating a mechanistic role for tau in epileptogenesis and network excitability. While previously viewed as a consequence of neurodegeneration, seizures and epileptiform activity are increasingly recognized as an accelerant of disease progression (25, 34-38). Consistent with these clinical observations, we recently demonstrated that seizures exacerbated tau spread and cognitive dysfunction in the five times familial AD (5XFAD) mouse model (18, 39). To further determine whether early periods of neuronal activity were involved in subsequent tau spread, we crossed the 5XFAD line with a genetic mouse model (targeted recombination in active populations; TRAP) to permanently label neurons that expressed cFos in response to seizure induction, and found heterogenous responses in the neurons that developed neurofibrillary tangle pathology.

The neurons that are activated by seizures show increased vulnerability to tau pathology compared to the surrounding populations in 5XFAD-TRAP mice (18). However, whether these populations contribute to tau propagation in a model of tauopathy are unknown.

Here, we used the TRAP cross with the T40PL-GFP mice, which harbor the P301L *MAPT* mutation labeled with GFP (T40PL-TRAP), to determine the contribution of activated neuronal networks and populations to the development of tau pathology, with the hypothesis that seizures would worsen tau propagation through activated neuronal networks and populations in the

T40PL-TRAP model, consistent with our observations in the 5XFAD model (18, 39). Using AD-tau seeding, PTZ seizure kindling, and brain-wide light sheet fluorescence microscopy (LSFM), we found that T40PL-GFP mice have increased seizure susceptibility compared to WT mice, and that seizure induction increased activity-dependent neuronal labeling and elevated tau-GFP levels preferentially in seizure activated neurons, indicating that this neuronal subpopulation disproportionately contribute to tau propagation in T40PL-TRAP mice. Our study supports a common mechanism for seizure-induced disease progression in tauopathies, AD, and possibly other related neurodegenerative diseases, which in turn may serve to identify promising therapeutic targets to slow tauopathy progression.

## Methods

### Mice

All experiments involving mice were approved by the University of Pennsylvania Institutional Animal Care and Use Committee (IACUC) Office of Animal Welfare. T40PL-GFP were first generated as previously described (17) and express the 2N4R human tau T40 isoform with P301L mutation and GFP tag driven by the mouse prion promoter. Heterozygote T40PL-GFP mice and littermate WT mice were used in PTZ kindling experiments. Fos^2A-iCreER^ (TRAP2) and B6.Cg-*Gt(ROSA)26Sor*^tm14(CAG-tdTomato)Hze^/J (Ai14) were purchased from Jackson Laboratory (Bar Harbor, ME). TRAP2 and Ai14 mice were bred for two generations to produce mice homozygous for both transgenes. Heterozygous T40PL-GFP mice were bred with TRAP2 x Ai14 mice to produce the T40PL -TRAP and WT-TRAP littermate mice that were used in the brain mapping studies. These mice contain a tamoxifen inducible, cFos driven Cre, which results in permanent expression of tdTomato in activated (cFos expressing) neurons in the 4-6 hours following 4-OHT administration (40).

Male and female mice were used for all experiments. Mice were randomly assigned to experimental groups and balanced for sex. Experimenters were blind to condition of mice during data acquisition and analysis.

### PTZ kindling

At 3 months of age T40PL-GFP mice underwent an established PTZ kindling protocol or control (Saline) procedures(39, 41). Briefly, mice were injected (I.P.) with 35 mg/kg PTZ (Sigma Alrich, St Louis, MO) or vehicle (0.9% saline, Sigma Aldrich) every 48 hours for a total of 8 injections. Mice were video recorded for 45 minutes and assessed for seizure severity with a modified Racine score (42): 0=normal behavior, 1=freezing, 2=hunching with facial automatism, 3=rearing and forelimb clonus, 4=rearing with forearm clonus and falling, 5=tonic-clonic seizure, 6=death.

### AD-tau isolation

Pathological tau protein was isolated from the cortex of patients with neuropathological AD diagnosis as previously established (15). BCA determined total protein concentrations and ELISA was used for tau (0.4-1.1 μg/μL), Aβ_42_ (32-132 ng/mL), Aβ40 (3-46 ng/mL), and α synuclein (nondetectable – 0.54 μg/mL).

### Stereotaxic Surgery

At 3 months of age, mice were deeply anesthetized with ketamine-xylazine-acepromazine and underwent stereotaxic surgery to inject AD-tau into the right hippocampus and overlying posterior parietal association cortex using previously established protocols (15-18). Briefly, a hole was bole -2.5 mm from Bregma and 2 mm from the midline. A Hamilton syringe was slowly lowered to a depth of 2.4 mm and 2.5 uL AD-tau (0.4 μg/uL) was injected over 2 minutes. The syringe was retracted to a depth of 1.4 mm and an additional 2.5 μL AD-tau (0.4 μg/uL) was injected over two minutes.

### 4-hydroxytamoxifen (4-OHT) administration

4-OHT (Sigma Aldrich) was dissolved with ethanol and suspended in an oil mixture of 4 parts sunflower oil (Sigma Aldrich) and 1 part castor oil (Sigma Aldrich). 4-OHT was administered via I.P. injection at 50 mg/kg on the final day of PTZ kindling for tamoxifen inducible, cFos driven permanent tdTomato labeling.

### Euthanasia

Cardiac perfusions with ice cold 4% paraformaldehyde took place following anesthesia with pentobarbital (I.P., 50 mg/kg) (Sagent Pharmaceuticals, Schaumburg, IL) for all experimental mice.

### Brain clearing, light sheet fluorescence microscopy (LSFM) and brain mapping

Brains used for LSFM and brain mapping were shipped to Life Canvas Technologies (Cambridge, MA) where they underwent brain clearing, antibody labeling, LSFM, and mapping. Briefly, SHIELD preservation was performed followed by SDS-based delipidation and refractive index homogenization (Clear and delipidation buffers (Life Canvas Technologies)) over 6 days (43). Samples were then immunolabeled using SmartLabel reagents (Life Canvas Technologies) (44) and 72 µg rabbit anti NeuN followed by incubation with secondary solutions (44). Intact, cleared, and immunolabeled brains were imaged using a SmartSPIM axially swept light sheet microscope (Life Canvas Technologies). 488 nm (tau-GFP), 561 nm (tdTomato), 647 nm (NeuN) lasers were used with a 3.6x objective and 4 μm Z-step. Resulting voxels were 1.8 μm × 1.8 μm × 4 μm.

Images were then aligned to the Allen Brain Atlas using an automated alignment process by Life Canvas Technologies. Brain-wide tau-GFP levels, and tdT+ and NeuN+ cells were mapped to the brain atlas. Atlas images in Fig 2 were generated using Mouse Brain Heatmap (45).

### Statistics

Racine scores between genotypes were compared by paired t test with Tukey’s post hoc. Tau-GFP and tdT+ mapping data were analyzed by hierarchical Bayesian modeling incorporating subregions nested within parent regions and the mouse sample as random effects. Rates of somatic tau-GFP were compared between tdT+ and tdT-neurons in PTZ treated T40PL-TRAP mice by Bayesian statistics incorporating cell type and brain region as fixed effects. For Bayesian statistics, credible effects were identified when credible intervals did not include 0.

Weakly informative priors were used based on pilot studies. Convergence, effective sample sizes and other details of Bayesian models can be found in Table S1. T tests were performed in Prism 10 (GraphPad, Boston, MA) and Bayesian statistics were performed in R (4.3.2).

## Results

### T40PL-GFP mice have increased seizure severity compared to WT mice

Given prior literature demonstrating increased seizure susceptibility in both Aβ (18, 39) and tauopathy rodent models (27, 28), we first examined the response of T40PL-GFP mice to PTZ kindling. To do so, we used an established PTZ kindling protocol involving injections of PTZ at an initially subconvulsive dose (35 mg/kg) every 48 hours over two weeks (8 total injections) and video-recorded animal behavior in the 45 minutes following PTZ administration and assessed seizure severity by modified Racine score (42). We found that T40PL-GFP mice have increased seizure severity overall compared to WT mice and a trend towards reduced latency for seizure onset (Fig 1). No mice died during PTZ kindling. These data demonstrate increased seizure severity and susceptibility in T40PL-GFP mice.

**Figure 1.**
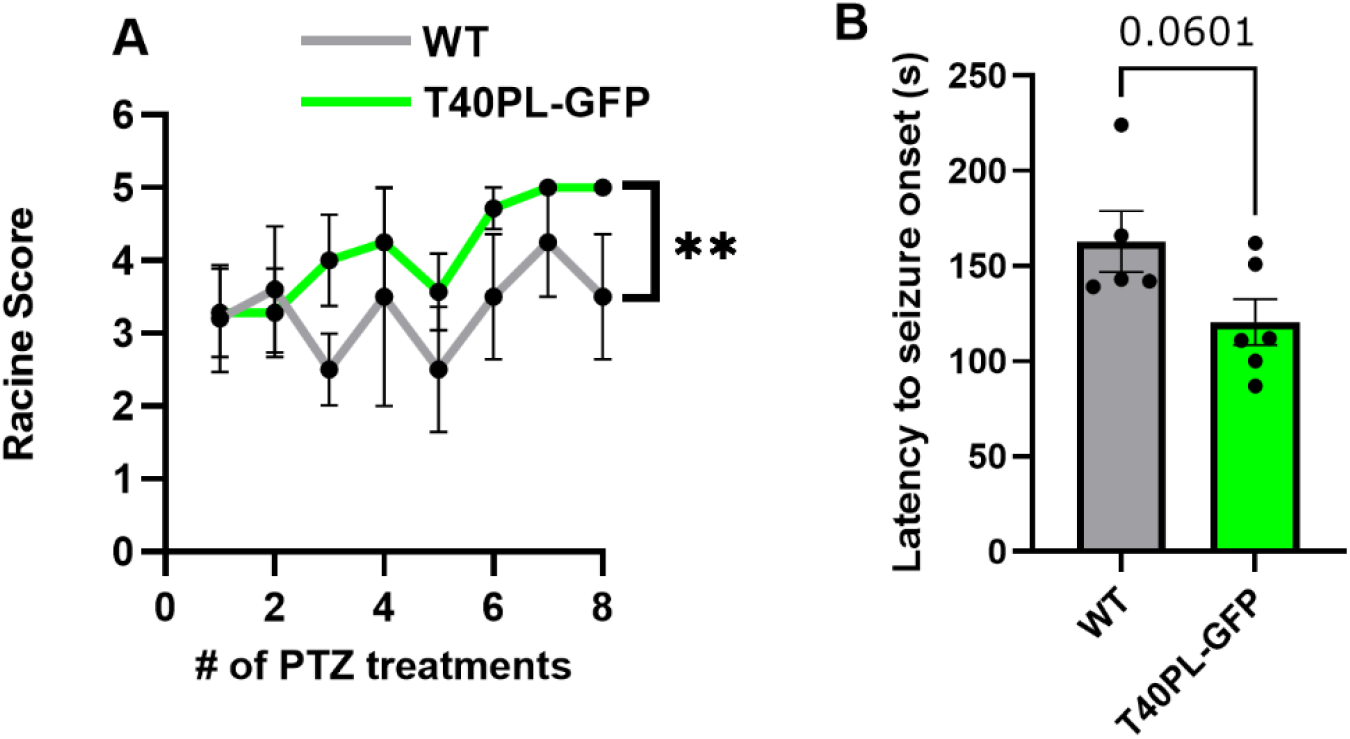
Increased seizure severity in T40PL-GFP mice during PTZ kindling. (A) A two-tailed t-test was performed to compare Racine scores across treatments between T40PL-GFP and WT mice. **=p<0.01, n=5-6/group. (B) Latency to seizure onset was compared by two-tailed t-test between T40PL-GFP and WT mice. WT = 5 mice (2 female), T40PL-GFP = 6 mice (3 female). Histograms display group mean ± standard error of the mean.

**Figure 2.**
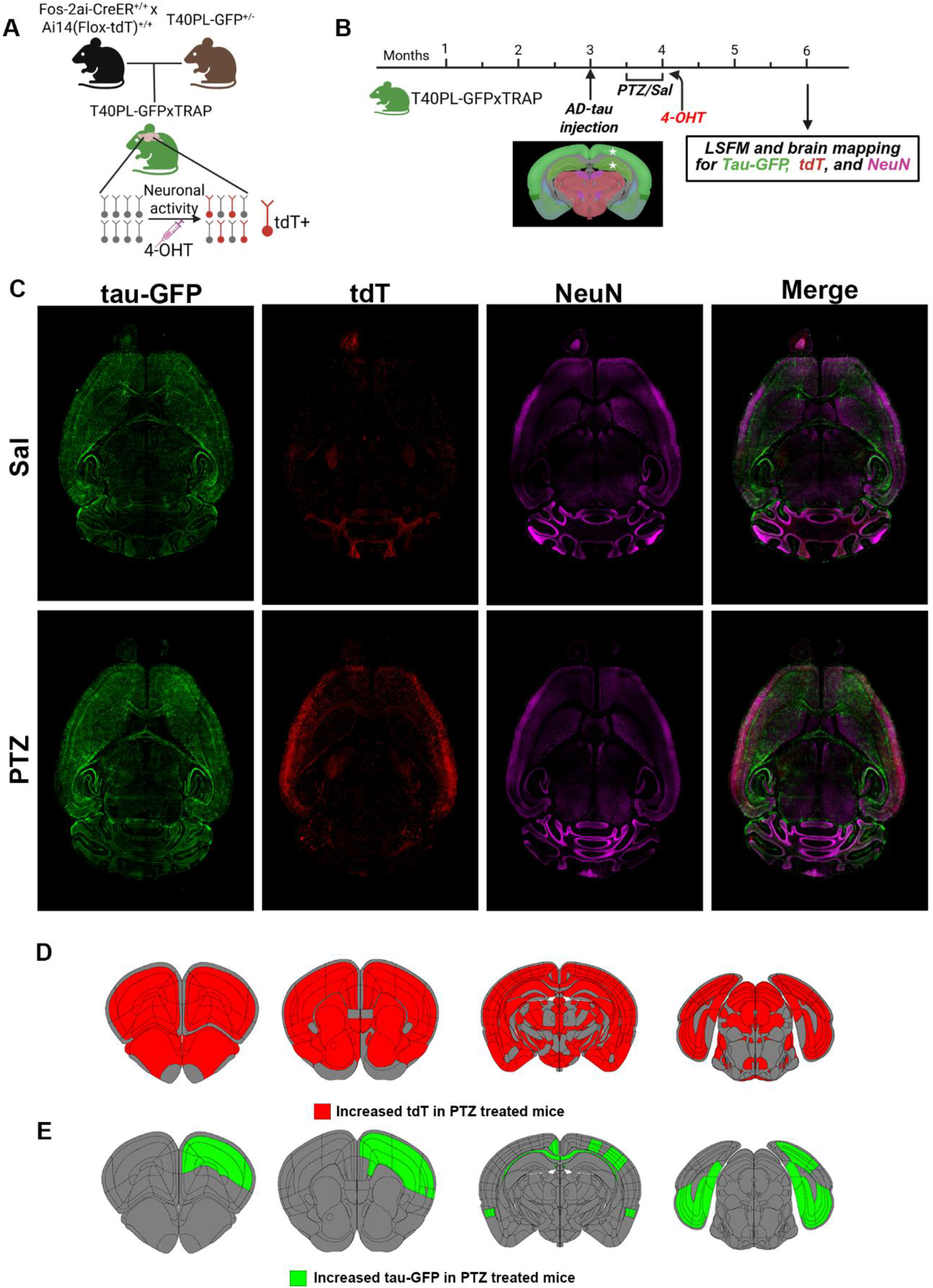
Seizures worsen tau-GFP and increases activity dependent labeling in T40PL-TRAP mice. (A) Schematic depicting the generation of T40PL-TRAP mice and 4-hydroxytamoxifen (4-OHT) induced, cFos driven tdT labeling. (B) Experimental timeline. (C) Representative 40 µm maximum intensity projections of horizontal images of Sal (top) and PTZ (bottom) treated T40PL-TRAP mice brains imaged by light sheet fluorescence microscopy. (D) Brain atlas schematics highlighting brain regions with credibly increased tdT levels in PTZ treated T40PL-TRAP mice compared to Sal treated T40PL-TRAP mice.(E) Brain atlas schematics highlighting brain regions with credibly increased tau-GFP levels in PTZ treated T40PL-TRAP mice compared to Sal treated T40PL-TRAP mice. n=4-5/group (2-3 female, 2 male). Details of Bayesian statistical models used identify credible changes in tau-GFP and tdT+ levels are included in Tables S2, S3.

### Brain-wide mapping reveals that seizures worsen tau pathology and activity dependent labeling in a novel tauopathy mouse model

Similar to our previous study in the 5XFAD mice (18), we generated a novel mouse model by crossing T40PL-GFP mice with TRAP mice to allow for permanent neuronal activity dependent labeling (Fig 1A). To promote tau propagation, we injected human AD brain derived tau lysate (AD-tau) into the right hippocampus and overlying cortex at 3 months of age (Fig 1B). T40PL-TRAP mice then underwent PTZ kindling (or control, Sal procedures) (18, 39) to induce seizures two weeks following AD-tau injection (Fig 1B). To examine the impacts of seizures on brain-wide tau pathology and activity dependent labeling we performed brain clearing, LSFM, and mapping of tau-GFP and tdT+ neurons to the Allen Brain Atlas.

To compare tdT+ and tau-GFP levels from brain-wide LSFM imaging between Sal and PTZ treated T40PL-TRAP mice, we used Bayesian statistical modeling as an approach robust to relatively low sample sizes and hierarchical data structures (brain subregion nested within parent structure), making it well-suited for these brain mapping data (18, 46). These models identify credible differences between treatment groups as opposed to statistical significance associated with frequentist approaches. To begin, we examined activity dependent labeling (tdT+ cell counts). As expected, we found elevated tdT+ levels in the majority of parent and subregions, including the hippocampus, thalamus, striatum, and cortex (Fig 1C, 1D, Table S2). These data corroborate our prior reports of increased activity dependent labeling due to seizure induction (18, 47).

We next examined brain wide tau-GFP levels using Bayesian statistics. We found credibly increased tau-GFP levels in PTZ treated T40PL-TRAP mice compared to WT-TRAP mice in several parent brain regions in the cortex, hippocampus, and white matter tracts (Fig 1C, 1E, Table S3). Specifically, we found credible increases in PTZ treated mice in the ipsilateral (to AD-tau injection site) somatosensory, motor, gustatory, visceral, visual, prelimbic and anterior cingulate cortical subregions, and corpus callosum, contralaterally in the retrosplenial cortex, and bilaterally in retrohippocampal regions (subiculum, entorhinal cortex) (Fig 1C, 1E, Table S3). Notably, all of these brain regions have known direct neuroanatomical connections to the injection site (right dorsal hippocampus and posterior parietal cortex) (48), suggesting that seizures enhance the propagation of tau pathology through the endogenous neuroanatomical connectome, which is further supported by our findings of elevated tau-GFP levels in the white matter of PTZ treated mice. Importantly, we found that all areas that showed increased tau-GFP levels also had elevated tdT+ levels, demonstrating a vulnerability of seizure activated networks to enhanced tau pathology. Together these data demonstrate that PTZ induced seizures worsens tau propagation in T40PL-TRAP mice.

### Neuronal populations activated during seizures have increased rates of somatic tau pathology

We have previously demonstrated that seizure-activated neuronal populations are preferentially vulnerable to tau spread in the 5XFAD model (18). We next determined whether in the absence of amyloid pathology, tdT+ neurons are more likely to develop somatic tau pathology than surrounding tdT- (NeuN+) neurons in regions with elevated activity dependent labeling in PTZ treated T40PL-TRAP mice (Fig 3, Table S4). To do so, we colocalized tau-GFP signal above background levels with tdT+ and tdT- (NeuN+) soma and calculated the proportion of each cell type with somatic tau pathology throughout all brain regions. Using Bayesian statistical modeling, we found that there were higher rates of somatic tau-GFP in tdT+ compared to tdT-neurons in the ipsilateral and contralateral striata, and in the contralateral anterior group of the dorsal thalamus in PTZ treated T40PL-TRAP mice (Fig 3, Table S4). Notably, these areas have direct neuroanatomical connections to regions with elevated overall tau-GFP in PTZ treated T40PL-TRAP mice (48). These results demonstrate that in PTZ treated T40PL-TRAP mice, early seizure activated neurons disproportionately contribute to tau pathology.

**Figure 3.**
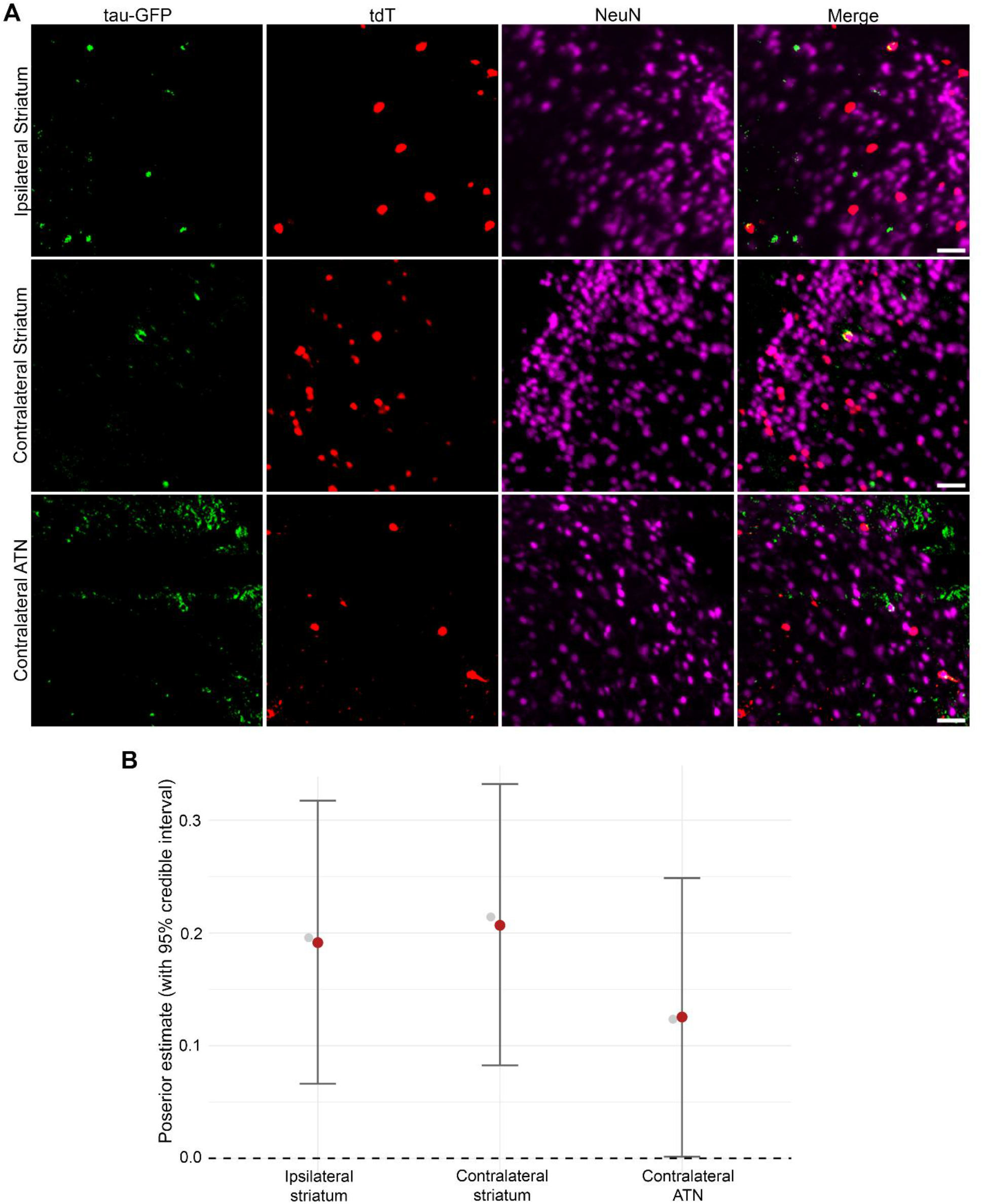
Increased rates of somatic tau-GFP in seizure activated neurons compared to surrounding neurons in PTZ treated T40PL-TRAP mice. (A) Representative light sheet microscopy images of tau-GFP, tdT, and NeuN from the ipsilateral (to AD-tau injection) striatum (top), contralateral striatum (middle) and contralateral anterior group of the dorsal thalamus (ATN) (bottom) from PTZ kindled T40PL-TRAP mice. Scale bar = 50 µm. (B) Forest plots displaying results from Bayesian statistical analysis where credible effects were found. Posterior estimates (red points) and credible intervals (black bars) of mean differences between the proportion of tdT+ and tdT- neurons with somatic tau pathology. Grey points indicate the raw differences between the proportion of tdT+ and tdT- neurons with tau pathology. N=4 mice (2 female, 2 male). Details of Bayesian statistical models used identify credible changes in somatic tau-GFP levels between tdT+ and tdT- neurons are included in Table S4.

## Discussion

Our study provides the first evidence that seizures worsen tau propagation in a tauopathy mouse model. We also found that PTZ elevated activity dependent labeling throughout the brain, supporting our prior results (18, 47), and that elevated tau-GFP was preferentially found in brain regions with increased activation in response to PTZ in AD-tau seeded T40PL-GFP mice. Furthermore, within PTZ treated mice, tdT+ neurons that were labeled following seizure induction showed increased rates of somatic tau-GFP compared to surrounding tdT-neurons. These data add mechanistic insight to clinical studies demonstrating accelerated neurodegeneration in patients with seizures (25, 34-38). Our study suggests that seizures and the neuronal networks and populations that are activated during seizures are promising targets to slow pathological tau progression.

TRAP models are widely used in basic research and we have recently crossed this line with 5XFAD mice (18). Here, we crossed TRAP mice with the T40PL-GFP mouse to generate a tauopathy model that allows for tamoxifen inducible, neuronal activity (cFos) dependent tdTomato labeling. Using the T40PL-TRAP mouse along with PTZ seizure kindling and AD-tau seeding, we explored our hypothesis that seizures worsen tau propagation in a tauopathy model.

We intentionally used PTZ as the chemoconvulsant because it induces seizures through hippocampal, cortical, and thalamic activation (18, 49, 50). Moreover, unlike other convulsants like kainate and pilocarpine, PTZ kindling results in minimal neuronal death in WT mice (18) (51, 52), making it a strong model to explore the impacts of seizures on neuronally driven tau spread without the confound of neuronal death. Our finding of worsened seizure severity in T40PL-GFP mice is consistent with prior reports using the Tau85/4 and THY-Tau22 mouse models (27, 28), further supporting a critical mechanistic role for tau in network excitability and seizure susceptibility. These findings are also in agreement with studies demonstrating that conditional and global tau knockouts reduce seizure susceptibility (29-33) even in the absence of tau pathology, supporting its therapeutic potential.

To comprehensively examine the impact of seizures on tau pathology we performed brain clearing and light sheet fluorescence microscopy to quantify and map brain-wide levels of tau-GFP and tdT+ neurons. While prior groups have used similar approaches to examine brain-wide tau pathology in rodent models (53, 54), these studies were largely descriptive. Here we used this approach to compare network-level tau propagation and activity dependent labeling between seizure induced and control mice. LSFM brain mapping datasets are hierarchical in nature, with brain regions nested within parent structures. This framework presents substantial statistical challenges, particularly when making comparisons between groups, with respect to multiple comparisons and limited sample sizes. Here we used Bayesian statistical modeling which uses partial pooling across regions, accounts for nested data structures, and provides group effect estimates while reducing false discovery inflation, making it well-suited to analyze these data (18, 46, 55). These features provide a robust framework through which we investigated the impacts of seizures on tau propagation and activity dependent labeling in T40PL-TRAP mice.

AD-tau seeding recapitulates the connectome-directed spread of tau pathology in both WT and transgenic mouse models (15, 17, 18). We injected AD-tau into the right hippocampus and overlying cortex of T40PL-TRAP mice to accelerate tau pathology and induce the development and spread of insoluble tau aggregates (17). T40PL-GFP mice develop tau-GFP mislocalized to neuronal somatodendritic domains, but do not spontaneously develop insoluble pathological tau species (17). While we did not directly distinguish endogenous tau-GFP from seeding induced aggregates of human tau, the brain clearing methodology used here included SDS-based delipidation (43). Thus, the measured tau-GFP signal following brain clearing is likely caused by seeding induced detergent-insoluble pathological tau aggregates likely composed of both mutant human and mouse tau (17). Prior studies in T40PL-GFP mice demonstrated that AD-tau seeding induces the spread of insoluble tau aggregates from the injection site to neuroanatomically connected brain regions (17). Similar propagation patterns were found following seeding in WT mice as well as in amyloidosis and other tauopathy mouse models, with accelerated spread found in the transgenic models (16, 18, 56-58). Here we are the first to explore the impact of seizures in tau propagation in a tauopathy model. We found that seizures increased tau-GFP levels to brain regions with direct neuroanatomical connections to the dorsal hippocampus, including the prefrontal and retrosplenial cortices, and in the fiber tracts. These regions overlapped with seizure-activated neuronal networks (increased tdT+ counts during PTZ administration), demonstrating a preferential vulnerability of these hyperactivated networks to tau propagation. These patterns of seizure exacerbated tau spread found here are consistent with those from our prior report in 5XFAD-TRAP mice (18), yet occurred in the absence of amyloid pathology, with elevations found in the cortex and fiber tracts. Increased tau pathology due to seizure induction in white matter suggests that seizures promote the axonal transport of tau, possibly leading to its spread in interconnected brain regions. While prior studies have shown increased seizure susceptibility in tauopathy models (27, 28), ours is the first to demonstrate that seizure induction worsens tau pathology in these mice. Overall, these data demonstrate that seizures promote enhanced tau pathology propagation through neuronal projections and seizure-activated networks in T40PL-TRAP mice.

We next determined the relative contribution of tdT+ and tdT-neurons to tau propagation throughout the brains of PTZ treated T40PL-TRAP mice. We found that tdT+ neurons were more likely to develop somatic tau-GFP compared to the surrounding tdT-populations in the striatum bilaterally, and contralateral anterior thalamic subregions. Notably, these are brain regions which receive direct inputs from the prefrontal cortex and retrosplenial cortex (anterior group of the dorsal thalamus) and motor cortex (striatum) (48), regions in which we observed elevated tau-GFP levels in PTZ treated relative to Sal treated T40PL-TRAP mice. These findings suggest that seizures worsened tau burden overall in these upstream brain regions, which then promoted the development of somatic tau-GFP preferentially in seizure activated neurons. It is plausible that these seizure-activated neurons may act as local seed sites, eventually leading to elevated overall tau pathology to this brain region. Future studies should examine earlier and later timepoints to determine whether tdT+ neurons in the prefrontal and retrosplenial cortices develop tau pathology earlier than neighboring tdT-populations, and at a later timepoint to determine whether the striatum and thalamus develop increased overall tau-GFP levels. This experimental framework using TRAP labeling and brain-wide imaging can be used for neuronal tracing between tdT+ populations with high tau pathology to identify whether tau is transmitted between these seizure-activated populations. Given the preferential vulnerability of these populations, future studies examining whether the functional manipulation of these neurons can slow tau spread and identification of the mechanistic targets that underlie their vulnerability to tau propagation are warranted. Prior evidence demonstrate that neuronal activity can induce the release of tau *in vivo* and promote the spread of tau pathology *in vitro*, supportive of a potential synaptic mechanism of tau spread. Given that seizure induction promotes long-term neuronal hyperexcitability (47), it is possible that elevated tau spread due to seizures may be caused by elevated synaptic activity. It is also plausible that seizure induction creates a positive feedback loop between neuronal hyperactivity and tau pathology in this model. Ongoing studies are investigating the potential role of neuronal activity in tau spread *in vivo*.

Overall, these studies further substantiate our recent study identifying seizures as a critical component of underlying the progression tau pathology in an amyloid environment (18) and demonstrate the vulnerability of seizure-activated neuronal networks and populations to tau propagation. Thus, seizure mitigation in tauopathies, including AD, should be prioritized in patient populations with these comorbidities.

## Acknowledgements

These studies were supported by funding from the National Institutes of Health Institute of Aging T32AG000255 (AJB) and R01AG077092 (DMT and FEJ), and the Alzheimer’s Association AARF-22-972333 (AJB).

We would like to thank Xiaofan Li and Sydney Zebrowitz for their help with stereotaxic surgery and PTZ kindling, respectively.

## Data availability

Data from these studies are available upon reasonable request from the corresponding authors.

## Notes

### Competing Interest Statement

The authors have declared no competing interest.

