## Supplemental material for "Seizures drive tau propagation in a tauopathy mouse model"

### Supplemental materials

<table>
<tr><th>Model</th><th>Rhat</th><th>Bulk ESS Range</th><th>Tail ESS Range</th></tr>
<tr><td>tdT+/mm^3~Treatment +(1+Treatment| ParentID/RegionID) + (1 | MsID)</td><td><1.01</td><td>3814 - 6393</td><td>5298 - 5432</td></tr>
<tr><td>tau-GFP~Treatment + (1 + Treatment | ParentID/RegionID) + (1 | MsID)</td><td><1.01</td><td>2462 - 10816</td><td>4499 - 11568</td></tr>
<tr><td>Proportion of cell type with somatic tau-GFP ~ Cell type * RegionID + (1 | MsID)</td><td><1.01</td><td>2247.772 - 23755.135</td><td>4133.283 - 10091.236</td></tr>
</table>

**Table S1. Bayesian model diagnostics.** Diagnostics from Bayesian statistical models for tdT+ levels (Fig 2D), tau-GFP levels (Fig 2E), and proportion of tdT+ and tdT- neurons with somatic tau-GFP (Fig 3). Rhat of less than 1.01 indicates strong Markov Chain Monte Carlo convergence. Bulk and tail effective sample size (ESS) indicates reliable estimates of median and tails of the posterior distribution, respectively.

**Table S2. Results from brain-wide tdT+ mapping.** Mean and standard error of the mean (sem) for tdT+/mm$^3$ for each brain region. Results from Bayesian hierarchical statistical model (tdT+/mm$^3$ ~ Treatment + (1 + Treatment | Parent ID/Region ID) + (1 | Mouse ID)) show the estimated relative change and 95% credible intervals (CI) associated with PTZ treatment in each brain region. Child level results were collapsed into their corresponding parent regions when credible effects were not found at the child level. Results were considered credible when 95% CI (L-95% CI, U-95% CI) does not cross 0. N=4-5 mice/group.

**Table S3. Results from brain-wide tau-GFP mapping**. Mean and standard of the mean (sem) for background subtracted tau-GFP fluorescence levels for each parent brain region. Results from Bayesian statistical models (tau-GFP ~ Treatment + (1 + Treatment | Parent ID/Region ID) + (1 | Mouse ID)) show the estimated relative change and 95% credible intervals (CI) associated with PTZ treatment in each brain region. Child level analyses were collapsed into parent results when credible effects were not found. All results displayed are at the parent level as no credible effects at the child level were found. Results are considered credible when 95% CI (L-95% CI, U-95% CI) does not cross 0. N=4-5 mice/group.

**Table S4. Results from brain-wide analysis of somatic tau-GFP in tdTomato+ and tdTomato- (NeuN+) neurons in PTZ kindled T40PL-TRAP mice.** Mean and standard error of the mean (sem) for the proportion of tdTomato (tdT)+ and tdT- neurons with somatic tau-GFP fluorescence levels above background. Results from Bayesian statistical model (proportion of cell type with somatic tau-GFP ~ Cell type * brain region + (1| Mouse ID)) show the estimated relative change and 95% confidence intervals (CI) associated with tdT labeling. Results are considered credible when 95% CI (L-95% CI, U-95% CI) does not cross 0. N=4 mice/group.
